# Exotic alleles contribute to heat tolerance in wheat under field conditions

**DOI:** 10.1101/2022.02.09.479695

**Authors:** Gemma Molero, Benedict Coombes, Ryan Joynson, Francisco Pinto, Francisco J. Piñera-Chávez, Carolina Rivera-Amado, Anthony Hall, Matthew P Reynolds

## Abstract

Global warming is one of the most significant threats to food security. With temperatures predicted to rise and extreme weather events becoming more common we must safeguard food production by developing crop varieties that are more tolerant to heat stress without compromising yield under favourable conditions. By evaluating 149 spring wheat lines in the field under yield potential and heat stressed conditions, we demonstrate how strategic integration of exotic material significantly increases yield under heat stress compared to elite lines, with no significant yield penalty under favourable conditions. Genome-wide association analysis revealed three marker trait associations, which together increase yield under heat stress by over 50% compared to lines without the advantageous alleles and was associated with approximately 2°C lower canopy temperature. We identified an *Aegilops tauschii* introgression underlying the most significant of these associations. By comparing overlapping recombination of this introgressed segment between lines, we identified a 1.49Mbp region of the introgression responsible for this association that increases yield under heat stress by 32.4%. The genes within this region were extracted from diverse *Ae. tauschii* genomes, revealing a novel *Ae. tauschii* MAPK gene, a SOC1 orthologue and a pair of type-B two-component response regulators. Incorporating these exotic alleles into breeding programmes could serve as a pre-emptive strategy to produce high yielding wheat cultivars that are resilient to the effects of future climate uncertainty with no yield penalty under favourable conditions.

## Introduction

Wheat is the most cultivated crop in the world with more than 216 million hectares grown annually (1), most of which is produced under temperate conditions (2). Heat stress is one of the major abiotic stressors that impacts global wheat production, reducing leaf area, crop duration and the efficiency of photosynthesis and respiration (3) as well as reducing floret fertility and individual grain weight (4). Together, these physiological consequences negatively impact productivity (3) with potential devastating effects. For example, in 2010 Russia saw a 30% reduction in wheat yield during their hottest summer in 130 years (5). Cases like this could become commonplace as global warming causes temperatures to rise and extreme weather events to become more frequent. Simulations predict that yields globally will fall by on average 6% for each 1°C increase in temperature (6), with some regions reaching 9.1% ± 5.4% per 1°C rise (7). Adaptation to future climate scenarios is vital to ensure global food security (8). Climatic instability, combined with environmental constraints, such as restricted supplies of irrigation water and arable land loss, emphasises the need for breeding strategies that deliver both increased yield potential during favourable cycles and resilience to abiotic stress and environmental constraints.

Such adaptation relies on genetic variation underlying the traits of interest; however, modern elite wheat material typically has limited genetic variation, particularly in the D genome (9), due to historic genetic bottlenecks (10, 11) compounded by intensive artificial selection by breeders (12). A strategy employed by CIMMYT to increase the genetic diversity of wheat pre-breeding material is to incorporate exotic parents in their germplasm via strategic crosses (11, 13). The most common exotic parents used are Mexican landraces (14) and primary synthetics, which are produced by hybridising tetraploid durum wheat with *Aegilops tauschii*, the ancestral donor of the D genome, to recreate hexaploid bread wheat (15); these synthetic lines act as a bridge to introduce durum and *Ae. tauschii* variation into modern hexaploid wheat. This approach has been successful in introducing disease resistance as well as drought and heat adaptive traits (16, 17). Landrace and synthetic material have been identified with superior biomass in comparison to elite lines under drought and heat conditions (18, 19) and elite lines that include landrace or synthetic material in their background have been developed in recent years for drought, heat, and yield potential conditions (20–22).

Challenges remain for the effective deployment of landrace and synthetic material. Only a small fraction of these vast collections of crop genetic resources have been evaluated for climate resilience traits and potential tradeoffs under favorable conditions have not been assessed. Currently, most of these genetic resources are unused (23) as breeders tend to avoid exotic materials because of large regions of poor recombination and a fear of linkage drag (24). Furthermore, despite evidence of the contribution of exotic material in wheat improvement, the physiological and genetic bases of heat tolerance in this material remain unclear.

Here, we evaluate a spring wheat panel in the field containing contrasting material controlled for phenology and plant height under heat stress and yield potential conditions. We explore yield and related physiological traits and compare exotic-derived lines with elite lines. We conduct a genome-wide association study to reveal genetic associations with heat tolerance traits and evaluate their impact under favorable conditions. Finally, we identify introgressed material overlapping a marker trait association (MTA) and employ a novel method downstream of GWAS, using *in silico* mapping to narrow down the interval, explore recombination and extract candidate genes from introgressed genomes.

## Results

### Physiological evaluation of HiBAP I under heat stress

To estimate the contribution of exotic material to heat tolerance and identify its genetic bases, we evaluated the High Biomass Association Panel I (HiBAP I) for two consecutive years under yield potential and heat stressed irrigated conditions in NW-Mexico (**Table S1**). HiBAP I represents an unprecedented resource of genetic diversity (25). It contains 149 lines, some of which are elite while others contain exotic material from Mexican landraces, synthetics, and wild relatives (**Fig 1a, Table S2**). All lines have agronomically acceptable backgrounds and a restricted range of phenology and plant height (21) which allows traits of interest to be evaluated without confounding effects.

**Figure 1.**
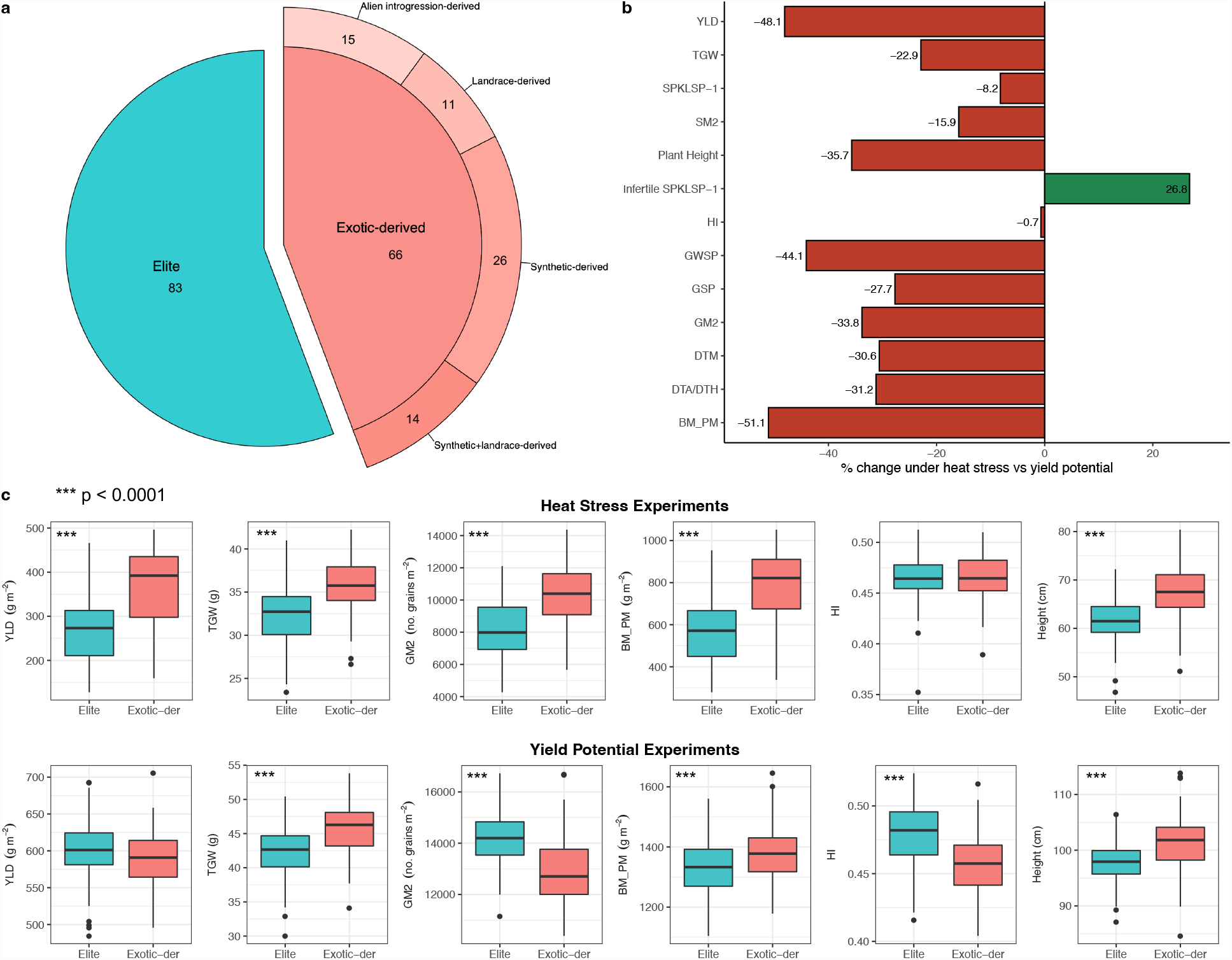
Physiological assessment of HiBAP I panel, comparing elite and exotic-derived lines under heat stressed and yield potential conditions. **a)** Number of lines from each group **b)** Effect of heat stress on traits, showing the percentage difference compared to yield potential conditions. **c)** Comparison of yield (YLD), thousand grain weight (TGW), grain number (GM2), biomass at physiological maturity (BM_PM), harvest index (HI) and plant height (PH) between elite and exotic-derived lines in HiBAP I measured under both heat stress and yield potential conditions. The significance of the differences between elite and exotic-derived lines for each trait was assessed using two-tailed t tests with no assumption of equal variance. *p*-values below 0.01 were considered significant (*), below 0.001 very significant (**) and below 0.0001 highly significant (***).

Heat stress was imposed by delayed sowing compared to the check environment (**Figure S1**) and, across both years of evaluation, this reduced yield by 48.1% and shortened the crop cycle duration by more than 30% (**Fig. 1b, Table S3**). When we analysed the response to heat stress of the lines based on their pedigree, exotic-derived lines exhibited an average of 37.7% higher yield compared to elite lines under heat stressed conditions (**Fig. 1c, upper**). Biomass, the trait most affected by heat stress, was 39% higher in exotic-derived lines, and other yield components, except for harvest index (HI), were significantly higher in exotic-derived lines than elite lines (**Fig. 1c, upper**). Under yield potential conditions, exotic-derived lines did not show a yield penalty compared to elite lines, as was reported by (**Fig. 1c, lower**). Exotic lines were on average 5.6cm and 3.8cm taller than elite lines under heat stressed and yield potential conditions, respectively. No differences in phenology were observed between the groups in either of the environments. Plant height was not correlated with yield under yield potential conditions (*r* = −0.007), but positive correlations were observed between plant height and yield under heat stressed environments (*r* = 0.699). The better performance of exotic-derived lines was validated using stress susceptibility index (SSI). This measure is negatively correlated with yield under heat stressed conditions; thus, lower SSI values indicate higher tolerance to a stressful environment. Compared to elite lines, exotic-derived lines had significantly lower SSI values for yield, grains per m^2^ (GM2) and biomass at physiological maturity (BM_PM), but not for thousand grain weight (TGW) (**Table 1**).

**Table 1.** Stress susceptibility index (SSI) calculated for yield (YLD), thousand grain weight (TGW), grains per m^2^ (GM2) and biomass at physiological maturity (BM_PM) of elite and exotic-derived lines obtained from adjusted means for two years of data in each environment. *r*_*p*_ corresponds to the phenotypic correlation with the yield obtained under heat environments. Data represents the mean ± S.D.

| Trait | $r_p$ (YLD_Heat) | Elite | | Exotic | |
| --- | --- | --- | --- | --- | --- |
| | | $n = 83$ | | $n = 66$ | |
| SSI_YLD | -0.976 | 1.16 ± 0.24 | **a** | 0.79 ± 0.32 | **b** |
| SSI_TGW | -0.439 | 1.03 ± 0.2 | **a** | 0.95 ± 0.27 | **a** |
| SSI_GM2 | -0.951 | 1.26 ± 0.42 | **a** | 0.60 ± 0.52 | **b** |
| SSI_BM_PM | -0.946 | 1.13 ± 0.20 | **a** | 0.85 ± 0.23 | **b** |

Letters indicate the statistical significance between Elite and Exotic groups. Means followed by different letters are significantly different ( $p$ -value < 0.01) according to pairwise t tests.

Additional physiological traits were measured in the experiments to help understand the physiological basis of the superiority of the exotic-derived lines under heat stressed conditions. Exotic-derived lines had significantly higher normalised difference vegetative index (NDVI), a proxy for biomass, and significantly lower canopy temperature (CT) during both vegetative stage and grain filling stages under heat stressed conditions but not under yield potential conditions (**Fig. 2**). NDVI measured during vegetative and grain filling stages was positively correlated with yield (**Fig. 2**) while CT was negatively correlated with yield at both stages (**Fig. 2**). These correlations were present under heat stressed conditions but not under yield potential conditions. The correlations were steeper for exotic-derived lines than elite lines for both traits across both phenological stages, suggesting that NDVI and CT is having a higher impact on yield in exotic-derived lines compared with elite lines. Under heat stressed environments, both NDVI and CT presented similar correlations with BM_PM, GM2 and other yield components (**Table S4**) but no correlation was observed under yield potential. The SSI index calculated for yield was negatively correlated with agronomic and physiological traits except for CT, where positive correlations were observed indicating that more tolerant lines had consistently cooler canopies (**Table S4**).

**Figure 2.**
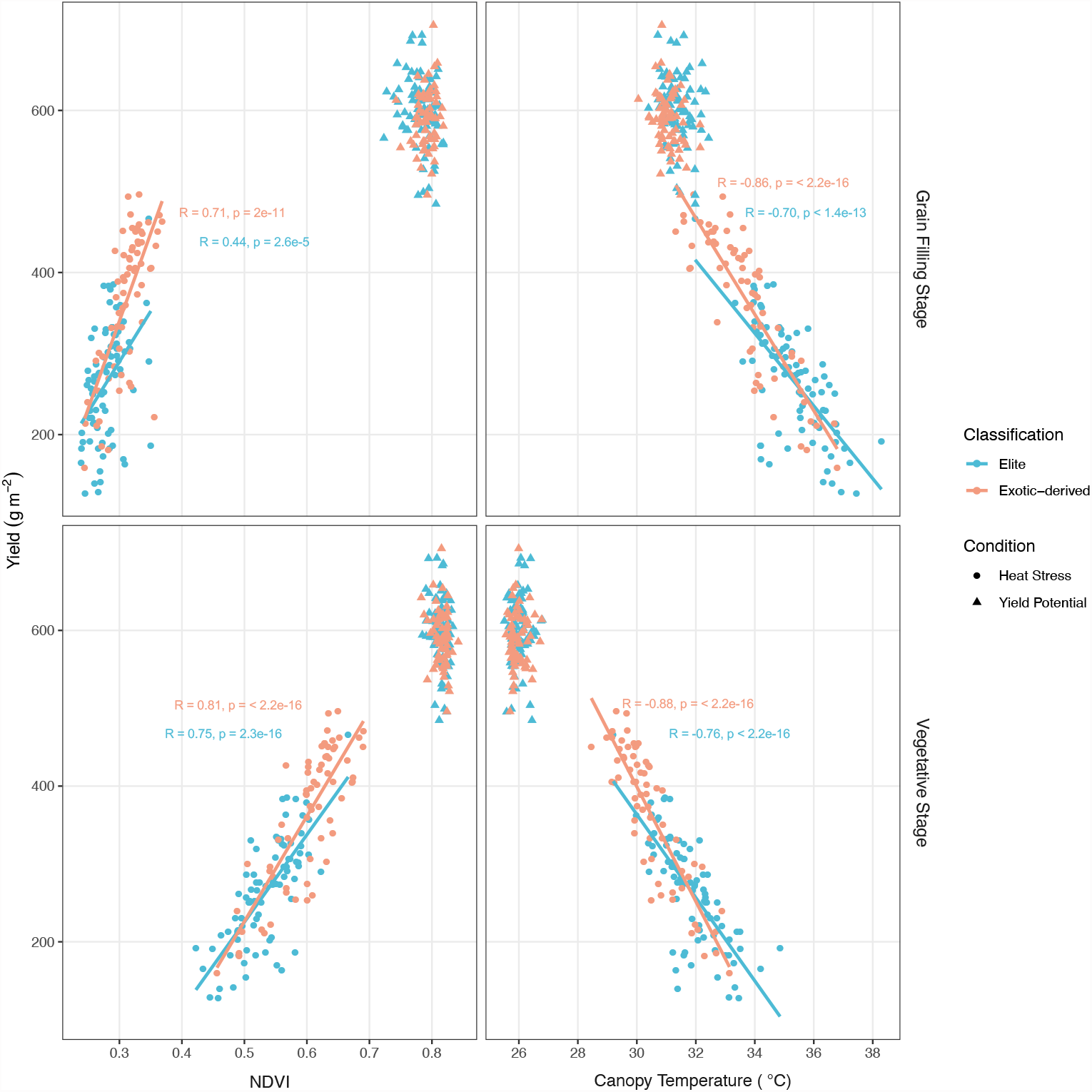
Relationship between normalized difference vegetation index (NDVI) and yield and between average canopy temperature (CT). NDVI and CT were measured with UAVs at pre-heading (vegetative stage) and during grain filling. Regression lines were added for groups with a significant correlation (*p-value* <= 0.01) calculated using Pearson’s correlation coefficient.

### Genome-wide association analysis reveals genetic associations under heat stress

To explore the genetic bases of these exotic-derived heat tolerance traits, marker-trait association analyses were performed using BLUE (Best Linear Unbiased Estimator) means from two or four replicates for each measured trait over two growing seasons. The most relevant MTAs are shown in **Table S5**, and all Manhattan plots are shown in **Figure S2**. We found 3 pleiotropic markers (**Fig. 3a**) on chr1B (chr1B-63398861: C), chr2B (chr2B-820002: C) and chr6D (chr6D-6276646: T). These MTAs were associated with all three heat stress indices along with multiple yield traits, including yield and canopy temperature, at both vegetative and grain filling stages, and were not associated with HI or phenology (**Fig. 3c, Fig S2)**.

**Figure 3.**
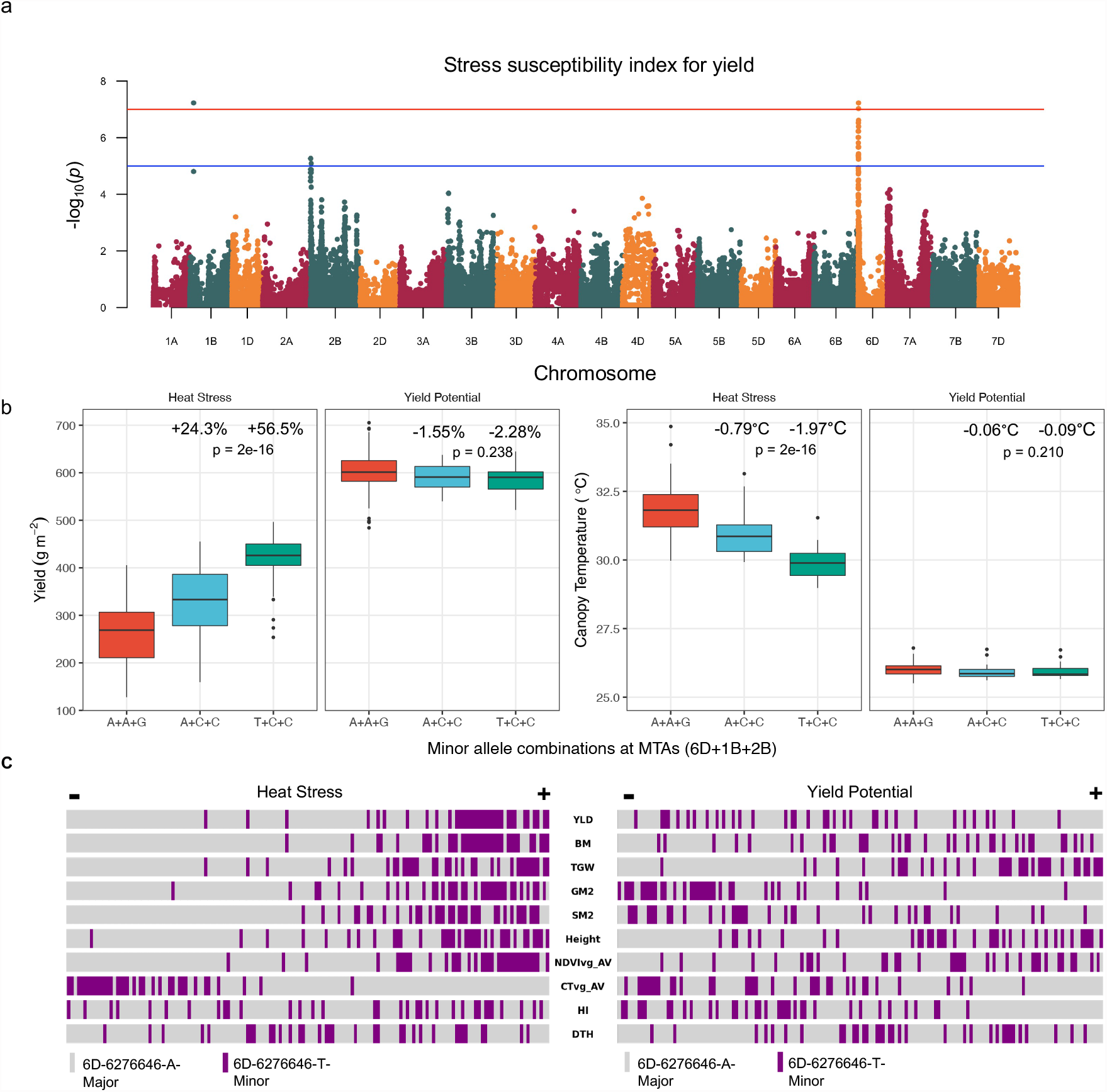
Genome-wide association study reveals genetic markers underlying heat tolerance traits. **a)** Manhattan plot showing the marker trait association for stress susceptibility index (SSI) for yield **b)** Specific marker allelic variants effects on yield and on canopy temperature under heat stress in chromosomes 6D (chr6D-6276646: T), 1B (chr1B-63398861: C) and 2B (chr2B-820002). The percentage change and °C change is calculated compared to lines with the major alleles at all three MTAs. Significance of allele combinations was computed using a one-way ANOVA test **c)** Phenotype distribution under heat stress and yield potential conditions highlighting the rank of 6D minor allele carriers for each phenotype where lines in the panel are ordered from lowest to highest value for each trait.

Lines with the minor C allele on 1B and 2B and the major A allele on 6D have 24.3% higher yield under heat stress; lines which also have the minor T allele on 6D have 56.7% higher yield under heat stress compared to lines with the three major alleles (**Fig. 3b**). Assuming the three alleles do not interact epistatically, the T allele on 6D can be estimated to increase yield under heat stress by 32.4%. Lines with the minor allele at all three MTAs show a reduction in canopy temperature of 1.97°C and 2.37°C, at vegetative and grain filling stages, respectively, when compared to lines with the major allele at all three positions (**Fig. 3b**). Under yield potential conditions, no difference was observed between minor and major allele combinations for yield or for canopy temperature **(Fig. 3b)**. The minor allele at each of these MTAs is predominantly found in exotic-derived lines with 50/55 (1B), 44/45 (2B) and 33/33 (6D) lines with the minor allele classified as exotic-derived (**Table S2**).

### Aegilops tauschii introgression underlies 6D MTA

Due to the better performance of exotic-derived lines under heat stress and exotic-derived lines possessing alleles for heat tolerance, we searched for introgressed material overlapping the MTAs. We detected introgressed material in HiBAP I lines by looking for regions with reduced mapping coverage and SNPs shared with *Ae. tauschii, Th. ponticum* or *S. cereale* but not with wheat cultivars Weebil, Pavon76 or Norin61. We identified introgressed *Ae. tauschii* material at the beginning of 6D in all 33 lines with the T/T genotype and all 7 lines with the A/T genotype. Multiple offspring of the same Sokoll Weebil1 cross show that recombination occurs readily within the segment, breaking up the segment into variable sizes (**Fig. S3**). The full-length, unbroken segment is 31.6Mbp in length, as seen in Sokoll (HiBAP_57) (**Fig. 4a**). By comparing the overlapping segments between lines, we found a 5.05Mbp and 6.85Mbp that is present in all lines with the T/T or A/T genotype at 6D-6276646 and absent in all the lines with the A/A genotype (**Fig. 4a**). In A/T lines, the introgression itself appears to be heterozygous, evidenced by intermediate mapping coverage deviation compared to the homozygous lines and by heterozygous SNPs matched to *Ae. tauschii*. Using chromosome and protein alignments, we anchored this core region to the *Ae. tauschii* reference genome, Aet v4.0 (26), and extracted the syntenic 1.49Mbp region between 4.63Mbp and 6.12Mbp that likely contains the gene(s) responsible for the MTA (**Fig. 4b, 4c**). We found no evidence of introgressed material overlapping the 1B or 2B MTAs.

**Figure 4.**
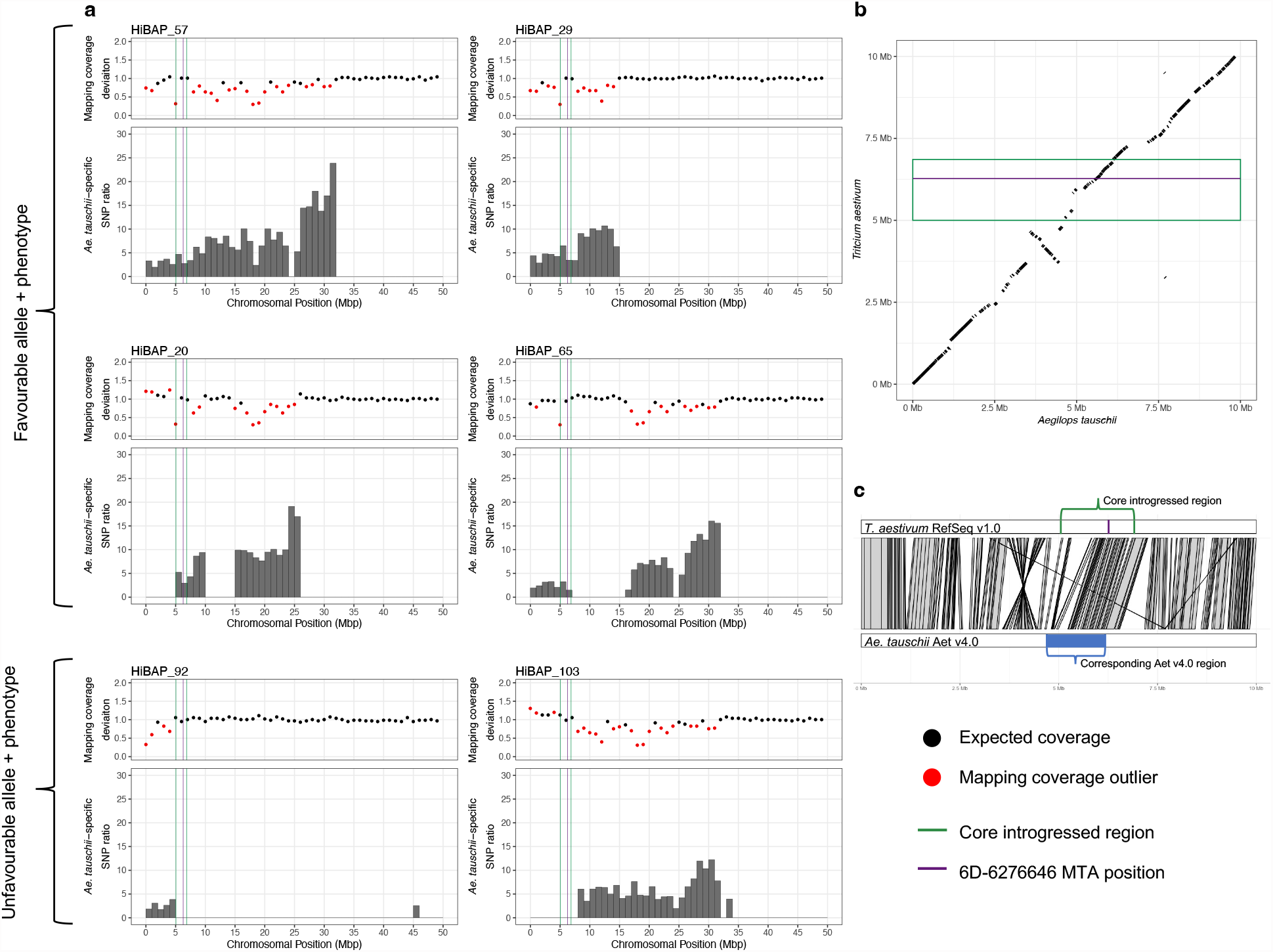
*Aegilops tauschii* introgression underlies chr6D-6276646 MTA. ***a*)** Visualising *Ae. tauschii* introgressions across the first 50Mbp of chr6D in six HiBAP I lines, four containing the favourable allele at chr6D-6276646 (HiBAP 57, 29, 48, and 65) and two containing the unfavourable A allele at chr6D-6276646 (HiBAP 92 and 103). Mapping coverage deviation was computed between the HiBAP line and the median of the panel in 1Mbp windows. Red points are statistically significant outliers. *Ae. tauschii*-specific SNP ratio in each 1Mbp window was calculated by dividing the number of homozygous *Ae. tauschii*-specific SNPs in that window by mean number of homozygous *Ae. tauschii*-specific SNPs in that window across the panel. Green lines mark the borders of the region common to all lines with the T genotype, corresponding to a 1.80Mbp region in CS and a 1.49Mbp in *Ae. Tauschii*. The purple line indicates the MTA position. ***b*)** Synteny between 6D:1-10,000,000 in CS RefSeq v1.0 (27) and *Ae. tauschii* Aet v4.0 (26). The green box indicates the 1.80Mbp region (1.49Mbp relative to *Ae. tauschii*) common to all lines with T genotype, corresponding to the green region in part a. The purple line indicates the MTA position **c)** Alignment of 6D:1-10,000,000 in CS RefSeq v1.0 (27) and *Ae. tauschii* Aet v4.0 (26), illustrating how the orthologous region in *Ae. tauschii* was identified and extracted.

### Candidate genes for MTAs in 1B, 2B and 6D

For the 6D MTA, we identified the syntenic region in the *Ae. tauschii* genome and a list of genes that had been introgressed. As we are unaware of the *Ae. tauschii* accession that has been introgressed, we also looked at the genes within the orthologous region in four other available chromosome-level *Ae. tauschii* assemblies (28). Between accessions, this region varies between 1.49Mbp and 1.82Mbp in length from and contains between 26 and 33 genes (**Table S6**). These include a MIKC-type MADS-box gene orthologous to SUPPRESSOR OF OVEREXPRESSION OF CONSTAN 1 (SOC1); a mitogen-activated protein kinase (MAPK) gene found in two of the *Ae. tauschii* accessions with no orthologue in wheat; and a pair of type-B two-component response regulator receiver proteins, orthologous to type-B *Arabidopsis* response regulators (ARRs) with closest similarity to ARR-11. One member of the pair, AET6Gv20025700, appeared to have a myb-binding domain that is missing from the wheat orthologue gene model. However, after manual reannotation, we found that this difference was a misannotation in wheat so likely not causing a functional difference. We also found that both ARR genes were expressed specifically in spike and grain in both *Ae. tauschii* and wheat, whereas the other candidate genes were expressed across leaf, root, spike and grain tissues. For the 1B and 2B MTAs, as they were not within an introgression, we submitted the sequence 1Mbp up and downstream of the MTA to Knetminer, a gene discovery tool (29). Within the 2B interval, we 10identified DEHYDRATION-RESPONSIVE ELEMENT-BINDING PROTEIN 1A (DREB1A) and STEROL GLUCOSYLTRANSFERASE (SGT) as promising candidate genes.

## Discussion

### Exotic-derived lines have higher heat tolerance with no yield penalty under favorable conditions

Exotic parents are routinely used to increase genetic diversity in wheat pre-breeding pipelines and their enhanced performance has been demonstrated under salinity (30), drought (14, 19) and heat stress (18, 31). In the present study, exotic-derived lines performed better under heat stress than elite lines with no yield penalty under yield potential conditions. This increased yield under heat stress was associated with a range of factors, including higher biomass throughout the crop cycle, higher grain number and cooler canopy temperature during both vegetative and grain filling stages. Cooler canopies have been previously associated with higher tolerance to drought and heat irrigated environments (32) and with optimised root distribution in bread wheat (33). Plants with an optimized root system are able to satisfy the high evaporative demand through elevated transpiration rates under hot irrigated conditions and thus maintain cooler canopies (34). Higher transpiration rates are associated with increased stomatal conductance that, in turn, is associated with higher photosynthesis that can explain the higher biomass observed in exotic-derived lines in comparison with elite lines. However, according to temperature response models in wheat (6), the observed reduction in plant temperature of approximately 2°C would be unlikely to account alone for the >50% increased yield of exotic lines according to temperature response models in wheat (6). Despite variation in plant height being restricted, exotic-derived lines were taller than elite lines in both environments. Taller plants have a better light interception and light distribution, in comparison with shorter genotypes, and this has been associated with increased photosynthesis (35). Therefore, plant height can also influence the better performance of exotic-derivatives. Among all stress indices, the stress susceptibility index (SSI) is thought to be the most useful index for evaluating tolerant cultivars. Exotic-derived lines had significantly lower SSI than elite lines, adding additional support to the resilience of this exotic material under heat stress.

### Combining high density genotyping and high-throughput phenotyping to uncover novel markers for heat stability

The value of *de novo* SNP discovery through sequencing in breeding efforts is starting to be more widely recognised. In conjunction with high throughput phenotyping methods (36), high density, unbiased markers can be leveraged to discover novel MTAs or to narrow pre-existing QTL intervals to provide more robust markers for global breeding programs (37, 38). Using these methods, we have identified three pleiotropic markers on chromosomes 1B, 2B and 6D that when stacked increase yield by 56.5% and reduce canopy temperature by 1.97°C/2.37°C under heat stress conditions when compared to lines containing the three major alleles at these positions (**Fig. 3b**). These markers were associated with multiple agronomically important traits under heat stress including yield, grain per square meter, grain filling rate and biomass (**Fig. S2**). Despite being in apparently disparate regions of the genome, the 1B and 2B minor alleles always occur together and the 6D minor allele usually occurs with the 1B and 2B minor alleles. This suggests that there may be functional linkage between the markers. All three MTAs are predominantly found in exotic-derived lines but are not exclusive to any of the exotic categories as we see them in synthetic, introgression line and landraces derivatives. This brings their origin into question as their most recent pedigree suggests that the favorable alleles have come from different sources.

### Aegilops tauschii introgression underlies heat tolerance MTA

By utilising mapping coverage information and species-specific SNPs, we identified that the MTA on 6D was within an *Ae. tauschii* introgression. We show that this introgression readily recombines within CIMMYT germplasm by comparing the introgressed segment in different offspring off the same cross. Due to concerns regarding linkage drag and lack of recombination of wild relative introgressions (38), this is promising for the deployment of introgressed segments from the primary genepool into breeding programmes. The recombination enabled us to reduce the size of the interval responsible for the MTA by looking for the region always present in lines with the favourable genotype. This is a novel technique to employ downstream of a GWAS, akin to hapmapping or *in silico* genetic mapping. The smallest segment in the panel that contains the MTA is around 5Mbp and can likely be broken down further; the small size and it’s telomeric location make it amenable for deployment in breeding programmes.

The longest unbroken segment is present in Sokoll, a commonly used advanced synthetic-derived line. Recombination within the segment takes place in all Sokoll × Weebil1 crosses yet is unbroken in Sokoll. Therefore, Sokoll may be the donor line for this marker in many of the lines in HiBAP I. This would make sense given its presence in many of the pedigree histories of CIMMYT’s synthetic derived lines (**Table S2**). Some of the HiBAP I lines contain an *Ae. tauschii* segment that contains both the 6D MTA and a resistance gene upstream that underlies an MTA from a recent GWAS in *Ae. tauschii* (39). If the accession of *Ae. tauschii* in the HiBAP I lines confers the same resistance these lines could be used as donors for both traits.

The 6D MTA uncovered is supported by an MTA for heat tolerance reported in (13, 22) nearby on 6D. Singh et al., 2018 (13) state that the 6D MTA overlapped with an *Ae. tauschii* introgression, using speculative markers and pedigree-based inference. Here, we confirm this speculation and then demonstrate its ability to recombine and narrow down the introgressed region conferring the heat resilient phenotype through *in silico* introgression mapping.

### Putative candidate genes underlying heat stability

Following the identification of the core introgressed region, we extracted the syntenic orthologous region from five *Ae. tauschii* chromosome-level assemblies and used these as our source for putative candidate genes rather than the wheat reference genome. When appropriate, using non-reference genomes is important because the variation underlying the trait of interest might be absent from the reference genome. To select candidate genes for the observed yield stability under heat stress, we carried out extensive literature searches on the introgressed *Ae. tauschii* genes or the wheat genes within the interval (for the 1B and 2B MTAs); this uncovered several new candidate genes for further dissection.

We found two type-B two-component response regulator genes, with members of this family acting as transcription factors in the cytokinin signalling pathway (40). They have been linked to photoperiod stress protection via root-derived cytokinins (41) and negative regulation of drought response (42) via root cytokinin pathways (43); the resulting elevated levels of cytokinin have been linked to heat stress tolerance (44). The orthologues do not have identical protein sequences which could underlie a functional difference. However, these genes are expressed only in spike and grain in Chinese Spring and *Ae. tauschii* (if this holds following introgression) which may make these genes unlikely to be involved in the heat tolerant phenotype as it is established during the vegetative stage and maintained through grain filling. We also found a MIKC-type MADS-box transcription factor orthologous to the *Arabidopsis* gene SOC1, overexpression of which leads to chloroplast biogenesis, elevated photosynthesis, and tolerance to prolonged heat stress (45). Despite the protein sequences of the *Ae. tauschii* and wheat orthologues being identical, introgressed genes are often expressed differently when placed in a different genomic background (46) and upregulation of this gene may lead to the heat tolerant phenotype seen in SOC1 overexpression lines (45). Finally, we identified a novel MAPK gene. MAPKs play important roles in regulating responses to abiotic stress. They have been linked to oxidative stress tolerance under heat stress in wheat (47) and are involved in the heat stress response in *Arabidopsis*, maize and rice (48). The novelty of this gene and the presence-absence variation between *Ae. tauschii* accessions suggests this gene is recently evolved and possibly involved in environmental adaptation. The gene DREB1A was identified within the chr2B-820002 interval; this gene is part of a family of plant-specific transcription factors that bind DRE/CRT elements in the response to abiotic stresses. Overexpression of DREB1 in wheat led to drought tolerance and increased photosynthetic efficiency (49) and DREB2 overexpression lines displayed cold and heat tolerance (50). The 2B interval also contained STEROL GLUCOSYLTRANSFERASE (SGT) which has been identified to affect heat stability in both knockout and overexpression studies in *Arabidopsis* (51, 52).

Our proposed candidate genes for the 6D MTA differ from those proposed by Singh et al., 2018 (13). By using the *Ae. tauschii* genomes rather than relying solely on the wheat reference genome, we have demonstrated that the isoflavone reductase gene they proposed is not present in the core introgressed region. The other candidates were possibly disregarded due to lack of non-synonymous mutations identified from their markers; however, this ignores the possibility of differences in expression underlying a functional difference. These differences highlight the value of considering non-reference genomes downstream of a GWAS, particularly when divergent material has been introduced and the phenotype may be non-allelic in nature.

### Implications for breeding for climate change

These three markers can be deployed into marker-assisted breeding or introgression pipeline programmes to incorporate heat resilience traits into elite cultivars. The fact that no yield penalty was identified under more favourable conditions adds value to their deployment, especially given the negative impact that has been documented in terms of yield stability under increasing temperatures using extensive international data (53). The donor lines for these markers will be selected using our introgression mapping approach to introduce minimal linkage drag alongside the traits of interest.

## Materials and Methods

### Plant material and Growth Conditions

The High Biomass Association Mapping Panel HiBAP I consisted of 149 spring wheat lines (**Table S2**) and is composed of elite high yielding lines and lines with exotic material in their pedigree history. These exotic lines include primary synthetic derivative lines; Mexican and other origin landraces derivatives and elites containing introgressions, such as from *Thinopyrum ponticum* and *Secale cereale*. All lines have agronomically acceptable backgrounds and a restricted range of maturity and height to avoid extreme phenology or height confounding the expression of biomass and other traits. HiBAP I was evaluated during 2015/16 and 2016/17 under yield potential (referred to hereafter as YP16 and YP17, respectively) and heat stressed conditions (Ht16 and Ht17). Heat stressed conditions were created with delayed sowing where emergence was registered in March instead of November or December as in a normal growing cycle (**Table S1, Fig. S1**).

The field experiments were carried out at IWYP-Hub (International Wheat Yield Partnership Phenotyping Platform) situated at CIMMYT’s Experimental Station in Campo experimental Norman E. Borlaug (CENEB) in the Yaqui Valley, near Ciudad Obregon, Sonora, Mexico (27°24’ N, 109°56’ W, 38 masl) under fully irrigated conditions for both yield potential and heat stressed experiments. The soil type at the experimental station is a coarse sandy clay, mixed montmorillonitic typic caliciorthid. It is low in organic matter and is slightly alkaline (pH 7.7) (54). Experimental design for all environments was an alpha-lattice. Yield potential experiments consisted of four replicates in raised beds (2 beds per plot each 0.8 m wide) with four (YP16) and two (YP17) rows per bed (0.1 m and 0.24 m between rows respectively) and 4 m long. For heat stressed experiments, two replicates were evaluated for HiBAP I in 2m×0.8m plots with three rows per bed (**Table S1**). Seeding rates were 102 Kg ha^−1^ and 94 Kg ha^−1^ for YP and Ht experiments, respectively. Appropriate weed disease and pest control were implemented to avoid yield limitations. Plots were fertilised with 50 kg N ha^−1^ (urea) and 50 kg P ha^−1^ at soil preparation, 50 kg N ha^−1^ with the first irrigation and another 150 kg N ha^−1^ with the second irrigation. Rainfall, radiation, maximum, minimum and mean temperature by month for all the years of evaluation are presented in **Fig. S1**.

### Agronomic measurements

Phenology of the plots was recorded during the cycle using the Zadoks growth scale (GS) (55), following the average phenology of the plot (when 50% of the shoots reached a certain developmental stage). The phenological stages recorded were heading for heat experiments (GS55, DTH), anthesis for yield potential experiments (GS65, DTA) and physiological maturity (GS87, DTM) for both experiments. Percentage of grain filling (PGF) was calculated as the number of days between anthesis and physiological maturity divided by DTM.

Plant height was measured as the length of five individual shoots per plot from the soil surface to the tip of the spike excluding the awns. Spike, awn and peduncle length were measured in five shoots per plot before physiological maturity (PM). Fertile (SPKLSP^-1^) and infertile spikelets per spike (InfSPKLSP^-1^) were also counted in five spikes per plot at PM.

At physiological maturity, grain yield and yield components were determined using standard protocols (56). Samples of 100 (YP16), 50 (YP17) or 30 (Ht16, Ht17) fertile shoots were taken from the harvested area at physiological maturity to estimate yield components. The sample was oven-dried, weighed and threshed to allow calculation of harvest index (HI), biomass at physiological maturity (BM_PM), spikes per square meter (SM2), grains per square meter (GM2), number of grains per spike (GSP) and grain weight per spike (GWSP). Grain yield was determined on a minimum of 3.2 m^2^ to a maximum of 4.8 m^2^ under yield potential experiments and 1.6 m^2^ under heat experiments. In yield potential experiments only, to avoid edge effects arising from border plants receiving more solar radiation, 50 cm of the plot edges were discarded before harvesting. From the harvest of each plot, a subsample of grains was weighed before and after drying (oven-dried to constant weight at 70°C for 48 h) and the ratio of dry to fresh weight was used to determine dry grain yield and thousand grain weight (TGW). GM2 was calculated as (Yield/TGW) × 1000. BM_PM was calculated from yield/HI. SM2 was calculated as BM_PM/(DW shoots/Shoot number).

### Unmanned Aerial Vehicle (UAV) for CT and NDVI estimation

Aerial measurements data for CT and NDVI was collected using different aerial platforms. Each year, the logistics and availability determined which platform could be used for measuring the heat trials. A summary of the platforms used, together with the cameras and the achieved resolutions, is presented in **Table S7**. The multispectral and thermal cameras were calibrated onsite by measuring over calibration panels placed on the ground before and after each mission. An exception were the aircraft missions, where a calibration performed at the airfield would not be representative of the trial conditions. The flights were designed as a regular grid of north-south flightpaths covering the whole trial with images that overlapped 75% in all directions to ensure a good reconstruction of the orthomosaic. The flights were performed under clear sky conditions at solar noon ±2 hours.

NDVI and CT orthomosaics were obtained from the aerial images using the software Pix4D (need to confirm details). The orthomosaics were then exported to ArcGIS where a grid of polygons representing each polygon was adjusted on top of the image. To avoid the border effect, the polygons were buffered 0.5 m from the north and south border of the plot. Finally, the pixel values were extracted using the ‘raster’ package in R. We extracted the value of all the pixels enclosed within each polygon and removed possible outliers and calculated the average per plot.

### Stress Tolerance Indices

To determine the effect of heat stress in the genotypes evaluated across years and panels, Stress susceptibility index (SSI) was calculated using data from yield potential (Yyp) and heat stressed (Yht) experiments. as:

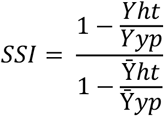

where 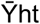 and 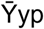 are the mean yields of wheat lines evaluated under heat stress and yield potential conditions, respectively (57).

### Statistical analysis for phenotypic traits

Data from both panels was analysed by using a mixed model for computing the least square means (LSMEANS) for each line across both years using the program Multi Environment Trial Analysis with R for Windows (METAR, (58)). When its effect was significant, DTA/DTH was used as a covariate (fixed effect) except for phenology. Broad sense heritability (*H*^2^) was estimated for each trait across both years as:

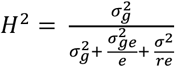

"where r = number of repetitions, e = number of environments (years), σ^2^ = error variance, σ^2^g = genotypic variance and σ^2^ge = G×Y variance. Unpaired *t-tests* for stress index (SSI) were conducted with the means across years to determine if the elite and exotic groups presented statistical differences with *p-value* <0.001.

### DNA extraction and Capture Enrichment and Genotyping

All genotyping data relating to the HiBAP I panel in this paper was taken from (25). Briefly, flag leaf material from 10 plants per line was collected from field grown plots and pooled prior to extraction. DNA was extracted using a standard Qiagen DNEasy extraction preparation and quality and quantity assessed. From this DNA, dual indexed Trueseq libraries with an average insert size of 450bp were produced for each line and enriched using a custom MyBaits 12Mbp (100,000 120bp RNA probe) enrichment capture synthesised by Arbor Bioscience. Prior to enrichment, samples were pooled into 8 samples per capture reaction. Post enrichment libraries were sequenced using an S4 flowcell on an Illumina NovaSeq S6000 producing 150bp paired end reads. These reads were mapped to the RefSeq v1.0 Chinese Spring reference genome (27) using BWA mem (59) and following filtering as per (25), SNPs were called with BCFtools (60) and subsequently filtered using GATK. SNPs were then annotated using SNPeff 4.3 (61). To create a set of shared SNPs for use in GWAS SNPs for all lines were combined and loci with more than 10% missing data and a minor allele frequency (MAF) below 5% were removed.

### Genome-Wide Association Study (GWAS)

GWAS analysis was conducted using the MLM method implemented in GAPIT (62). The effects of hidden familial relatedness were mitigated using principal component analysis eigenvectors 1-10 or membership coefficient matrices for 3-8 assumed subpopulations deduced by STRUCTURE (63) as covariates in the model. The EMMA (64) method was followed to create a kinship matrix required by the MLM method. Each MTA flanking interval was deduced by identifying the SNP position furthest upstream and downstream from the highest associated SNP that was above the −log P threshold of 5.

### Identifying regions of divergence

RefSeq v1.0 (27) was split into n genomic windows of size 1Mbp and 100Kbp using bedtools makewindows (65). Using the alignments produced in (25) and detailed above, the number of reads mapping to each window was computed using hts-nim-tools (66). To normalise by the sequencing depth of each line, read counts were divided by the number of mapped reads that passed the filters, producing normalised read counts c. Different windows of the genome have variable mapping coverage rates, so to compute coverage deviation we must compare each window to the same window in the other lines in the collection. Median normalised read counts, m, were produced, containing the median for each genomic window across the 149 lines. Mapping coverage deviation, d, was then defined for each line as:

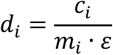

for window i ∈ {1, 2, …, n}, where ε is the median d value across the genome for the line. Statistically significant d values were calculated using the scores function from the R package ‘outliers’ with median absolute deviation (MAD) and probability of 0.99. This method was based on (67).

### Producing species-specific SNPs

Paired-end whole-genome sequencing data for the *Ae. tauschii* reference accession AL8/78 (26) and 5 additional accessions that represent 5 different clades (28), 4 *Secale cereale* accessions (68), *Secale. vavilovii* (68), *Thinopyrum ponticum* (69), and *T. aestivum* cultivars Weebil (69), Norin61 (69) and Pavon76 (46) were mapped to RefSeq v1.0 (27), filtered and SNP called as described for the genotyping above and in (25). Homozygous SNPs were retained if they had between 10 and 60 reads supporting the alternative allele and an allele frequency >= 0.8. Heterozygous SNPs were retained if they had between 10 and 60 reads mapped and were biallelic with each allele having >= 5 reads in support and an allele frequency >= 0.3. SNPs from one relative species not shared with any of the other species or wheat cultivars were retained as species-specific SNPs. These species-specific SNPs were assigned to HiBAP SNPs if they matched in position and allele. Species-specific SNP ratios were calculated by dividing the number of SNPs in each window matched to a species-specific SNP by the mean number of SNPs matched to that species in that window across HiBAP I. SNP ratio scores below 1.45 were removed to keep enriched scores only.

### Synteny between Ae. tauschii and T. aestivum

The first 10Mb of 6D from CS and *Ae. tauschii* Aet v4.0 (26) were aligned using Minimap2 (70) with parameters −x asm10. Alignments < 2.5Kb in length or with mapping quality < 40 were discarded. The dot plot was produced using pafr R package (71). Proteins encoded by genes in the first 10Mbp of 6D in *Ae. tauschii* and CS were aligned using BLASTp (72). Protein alignments and minimap2 alignments were used to anchor either side of the region commonly introgressed in all lines with the 6D T genotype to anchor the region from CS to *Ae. tauschii*. The *Ae. tauschii* genes and their proteins within this segment are considered as candidate genes. BLASTp (72) was used to compare these proteins to wheat proteins. Protein domains were identified using HMMER hmmscan (73) via ebi using Pfam, TIGRFAM, Gene3D, Superfamily, PIRSF, and TreeFam databases.

### Extracting corresponding region and genes from Ae. tauschii genomes

Proteins encoded by genes in the first 10Mbp of 6D in *Ae. tauschii* and CS were aligned using BLASTp (72). Protein alignments and minimap2 alignments were used to anchor either side of the region commonly introgressed in all lines with the 6D T haplotype to anchor the region from CS to *Ae. tauschii*. The sequence extracted from the *Ae. tauschii* reference genome was aligned to the other 4 chromosome-level assemblies using minimap2 (70) with parameters −x asm5. Alignments below length 5000 or quality of 40 were removed. The coordinates of each orthologous region were determined manually and the genes within these coordinates extracted from the respective gff files. The *Ae. tauschii* genes and their proteins within these segments were considered as candidate genes for functional exploration. BLASTp (72) was used to compare these proteins to wheat proteins. Protein domains were identified analysed using HMMER hmmscan (73) via ebi using Pfam, TIGRFAM, Gene3D, Superfamily, PIRSF, and TreeFam databases. Novelty of genes was determined by aligning the extracted protein sequence to each genome using tblastn (72).

### Exploring functionality of candidate genes

The genes in each identified interval (except for those in the 6D interval) were submitted to Knetminer (29). The knowledge networks created for each gene were then studied to identify links to the trait from which each MTA was deduced including their biochemical function and orthologous genes being linked in other organisms such as Rice and *Arabidopsis thaliana*. For the *Ae. tauschii* genes introgressed into the 6D interval, we conducted extensive literature searches to identify genes with links to heat stress response based on functional studies of related genes.

### Reannotating candidate gene and assessing tissue-specific expression

To test whether the missing myb-binding domain in the TraesCS6D02G014900 annotation was real or an artefact, we manually reannotated the gene. We identified the exon containing the MYB-binding domain in the wheat orthologue by aligning the coding sequence from the tauschii orthologue to Chinese Spring RefSeq v1.0 (27) using tblastn (72). We mapped Chinese Spring RNAseq data from leaf, root and shoot to RefSeq v1.0 (27) using HISAT2 (74) and assembled transcripts using cufflinks (75). We visually inspected the coding sequencing and RNA alignments IGV (76), which showed that the MYB-binding domain exon is present and expressed in wheat. To check whether the protein has a premature stop codon that might have interfered with the annotation and translation of the myb-binding domain, we extracted the coding sequence from the assembled transcript and checked for the presence of a complete open reading frame with no stop codons using EMBOSS getorf (77). Finally, we checked the presence of intact domains with HMMER hmmscan (73) via ebi using Pfam, TIGRFAM, Gene3D, Superfamily, PIRSF, and TreeFam databases. To explore qualitative expression of candidate genes, we mapped *Ae. tauschii* RNAseq data from leaf, root, seedling and developing grain 10dd (PRJEB23317) as above and abundances were counted using StringTie (78), taking the mean transcripts per million (TPM) across the replicates. Qualitative expression of the CS orthologues was explored using Wheat Expression Browser (79) and the previously leaf, root and shoot RNAseq data mapped above.

## Data availability

Publicly available sequencing data used in this study is available at the European Nucleotide Archive (ENA): *Th. ponticum* – SRR13484812; *S. vavilovii:* ERR505040, ERR505041, ERR505042; *S. cereale* accession Lo90: ERR504990, ERR504991, ERR504992; *S. cereale* accession Lo176: ERR505005, ERR505006, ERR505007; *S. cereale* accession Lo282: ERR505015, ERR505016, ERR505017; *S. cereale* accession Lo351: ERR505035, ERR505036, ERR505037; *Ae. Tauschii* accession XJ65: SRR13961980; Y173: SRR13962062; SX60: SRR13962012; AY29: SRR13961834; KU2832: SRR13961928; Y215: SRR13962048; Weebil1: PRJEB35709; Norin61: PRJNA492239; Pavon76:https://opendata.earlham.ac.uk/wheat/under_license/toronto/Hall_2021-10-08_wheatxmuticum/PIP-2495/200812_A00478_0126_AHN5W3DRXX/A10948_1_1/; *Ae. tauschii* RNAseq data: PRJEB23317; *T*.*aestivum* cv. Chinese Spring RNAseq data: Root - SRP133837; SRR6799264; SRR6799265; Leaf - SRR6799258; SRR6799259; SRR6799260; Spike - SRR6802608; SRR6802609; SRR6802610; SRR6802611.

Phenotypic data presented in this paper for the HIBAP I panel evaluated under yield potential and heat stressed environments can be found in the Dataverse CIMMYT Research Data Repository at https://data.cimmyt.org/dataset.xhtml?persistentId=hdl:11529/10548643

## Acknowledgements

This research was supported by the International Wheat Yield Partnership (IWYP) and by the Sustainable Modernization of Traditional Agriculture (MasAgro) an initiative from the Secretariat of Agriculture and Rural Development (SADER) and CIMMYT. Foundation for Food and Agriculture Research under the Grant ID: DFs-19-0000000013. AH was supported by BBSRC Core Strategic Programme Grant (Genomes to Food Security) BB/CSP1720/1; AH and RJ was supported by the BBSRC Designing Future Wheat grant BB/P016855/1, BBS/E/T/000PR9783 (DFW WP4 Data Access and Analysis). AH and RJ were also supported by BBSRC/IWYP BB/N020871/1. BC was supported by the BBSRC funded Norwich Research Park Biosciences Doctoral Training Partnership grant BB/M011216/1.

## Author contributions

AH, GM and MPR conceived of the idea and designed the experiment. GM, FP, FJPC and CRA collected the field data. GM analysed the physiological data. RJ conducted the genome-wide association study and Knetminer searches. BC conducted introgression analysis and introgressed candidate gene searches. BC, GM and RJ wrote the manuscript. AH and MPR were responsible for funding and supervision. All authors reviewed the manuscript.

## Competing interests

The authors declare no competing interests

## Notes

### Competing Interest Statement

The authors have declared no competing interest.

