## Supplementary figures for "Exotic alleles contribute to heat tolerance in wheat under field conditions"

**Figure S1**

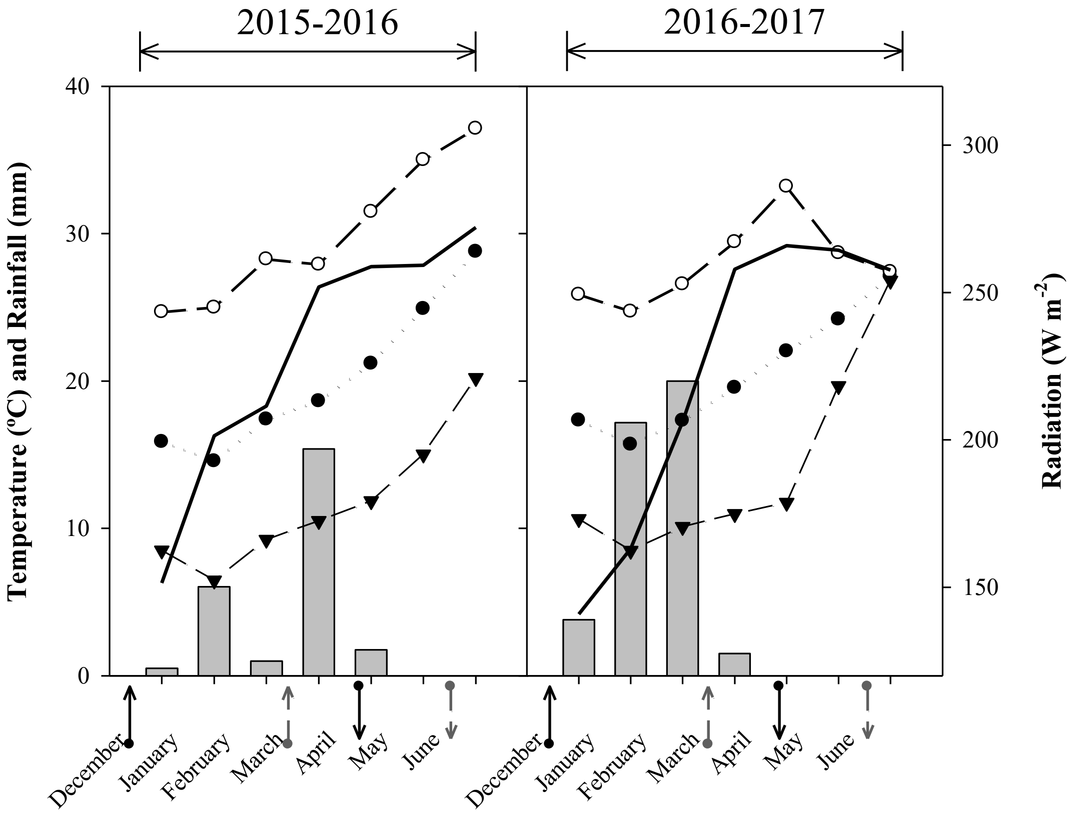


**Supplementary Figure 1.** Monthly accumulated rainfall (grey bars), average mean temperature (●), average maximum temperature (○), average minimum temperature (▼) and average monthly radiation (▬) registered during the experiments in the IWYP-HUB situated at CENEB experimental station situated in Ciudad Obregon Sonora, NW-Mexico. (↑) Represents emergence date for yield potential trials, (↓) represents harvest date for yield potential trials, (↑, broken line) represents emergence date for heat stressed trials, (↓, broken line) represents harvest date for heat stressed trials.

**Figure S2**

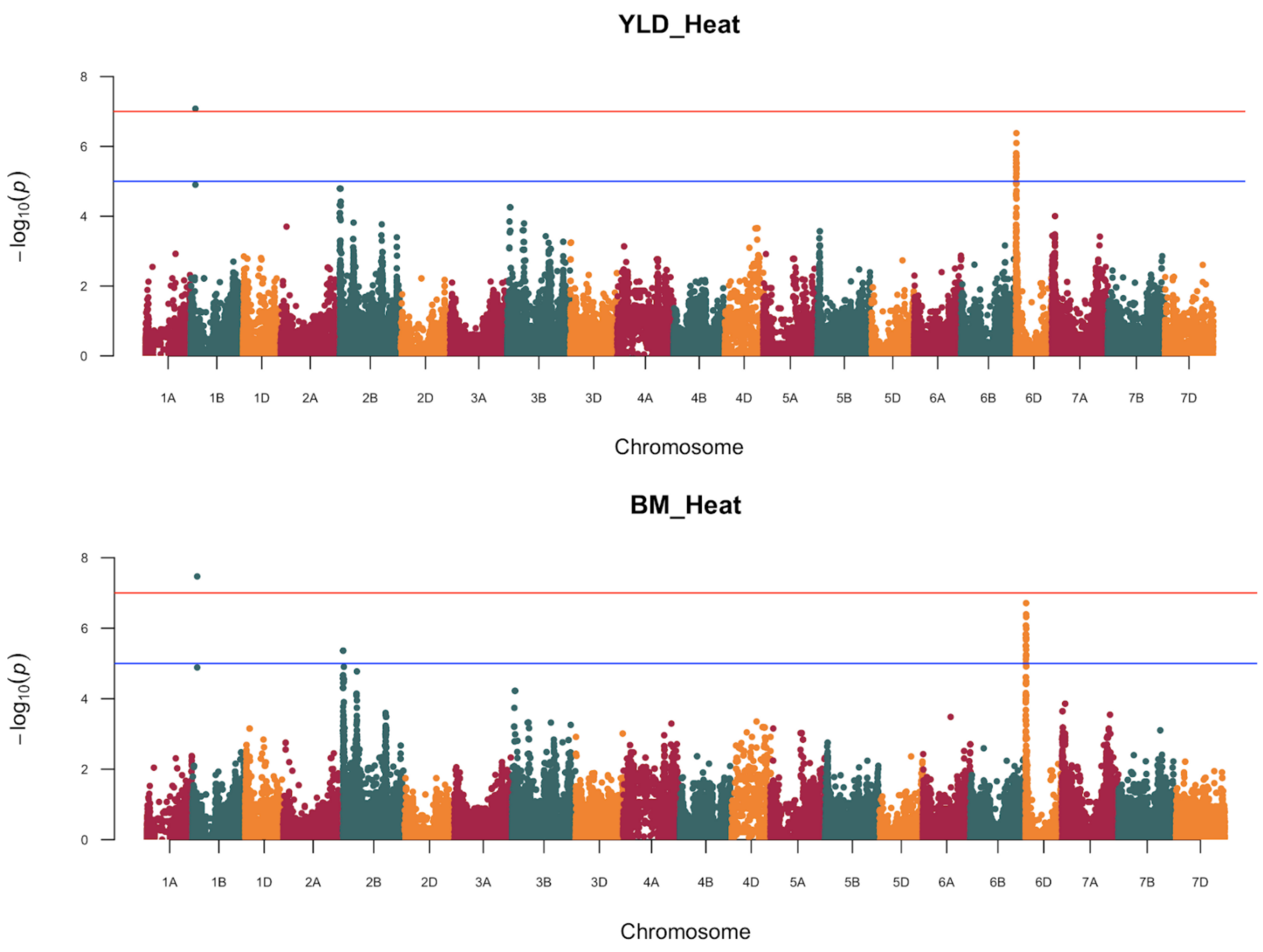
YLD_Heat
−log10(p)
8
6
4
2
0
1A
1B
1D
2A
2B
2D
3A
3B
3D
4A
4B
4D
5A
5B
5D
6A
6B
6D
7A
7B
7D
Chromosome
BM_Heat
−log10(p)
8
6
4
2
0
1A
1B
1D
2A
2B
2D
3A
3B
3D
4A
4B
4D
5A
5B
5D
6A
6B
6D
7A
7B
7D
Chromosome

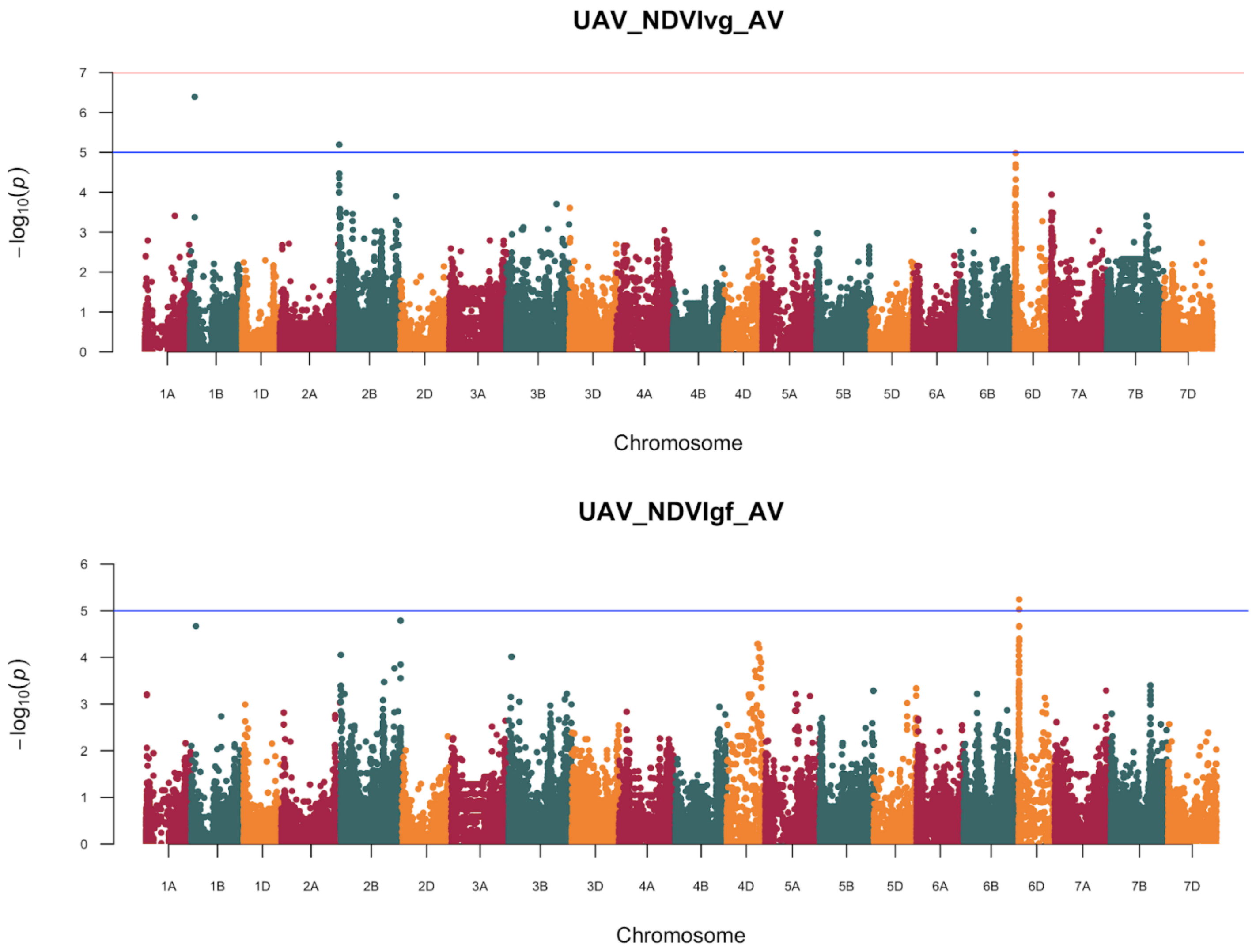
UAV_NDVIvg_AV
−log10(p)
0
1
2
3
4
5
6
7
1A
1B
1D
2A
2B
2D
3A
3B
3D
4A
4B
4D
5A
5B
5D
6A
6B
6D
7A
7B
7D
Chromosome
UAV_NDVIgf_AV
−log10(p)
0
1
2
3
4
5
6
1A
1B
1D
2A
2B
2D
3A
3B
3D
4A
4B
4D
5A
5B
5D
6A
6B
6D
7A
7B
7D
Chromosome

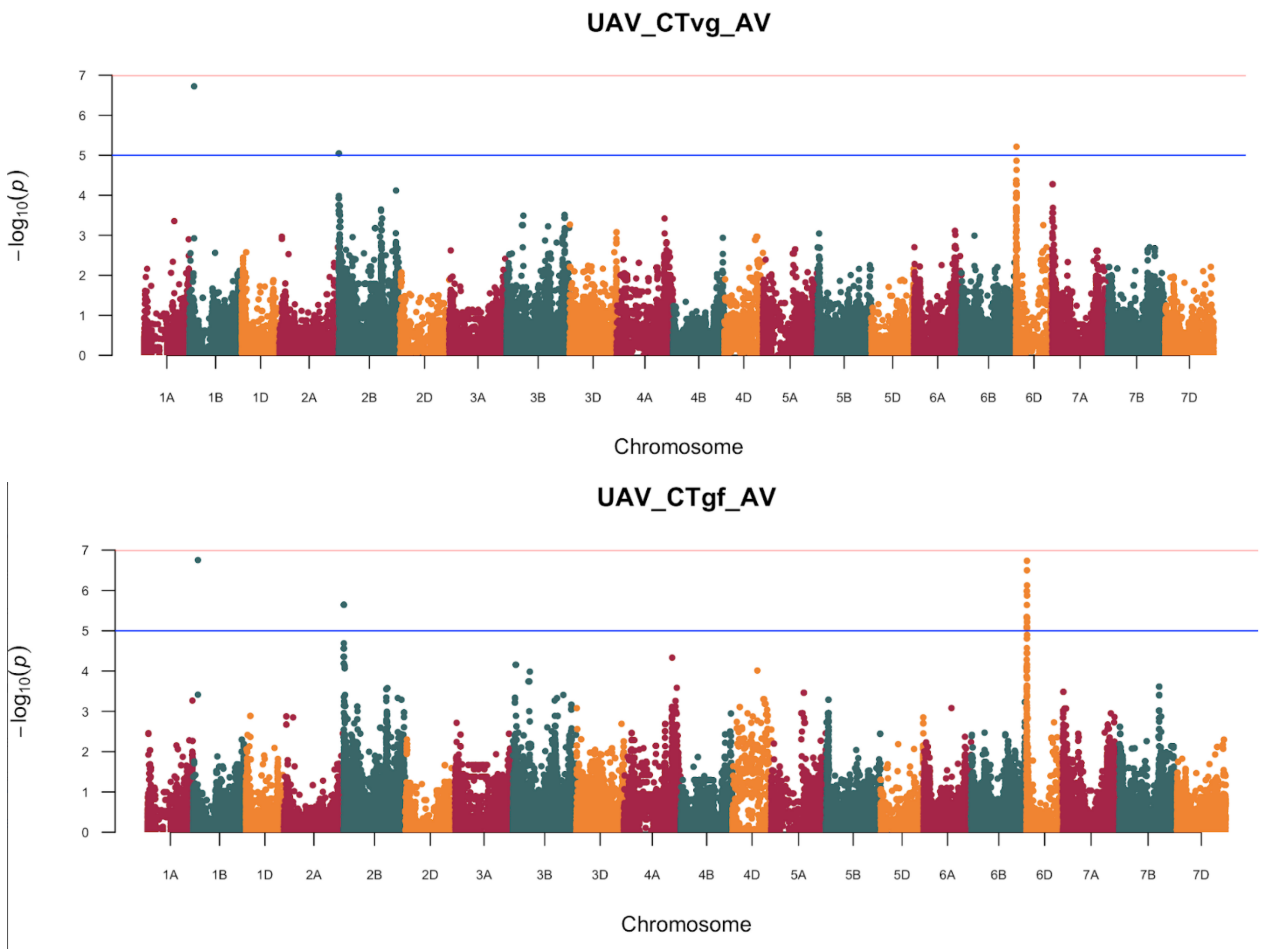
UAV_CTvg_AV
−log10(p)
0
1
2
3
4
5
6
7
1A
1B
1D
2A
2B
2D
3A
3B
3D
4A
4B
4D
5A
5B
5D
6A
6B
6D
7A
7B
7D
Chromosome
UAV_CTgf_AV
−log10(p)
0
1
2
3
4
5
6
7
1A
1B
1D
2A
2B
2D
3A
3B
3D
4A
4B
4D
5A
5B
5D
6A
6B
6D
7A
7B
7D
Chromosome

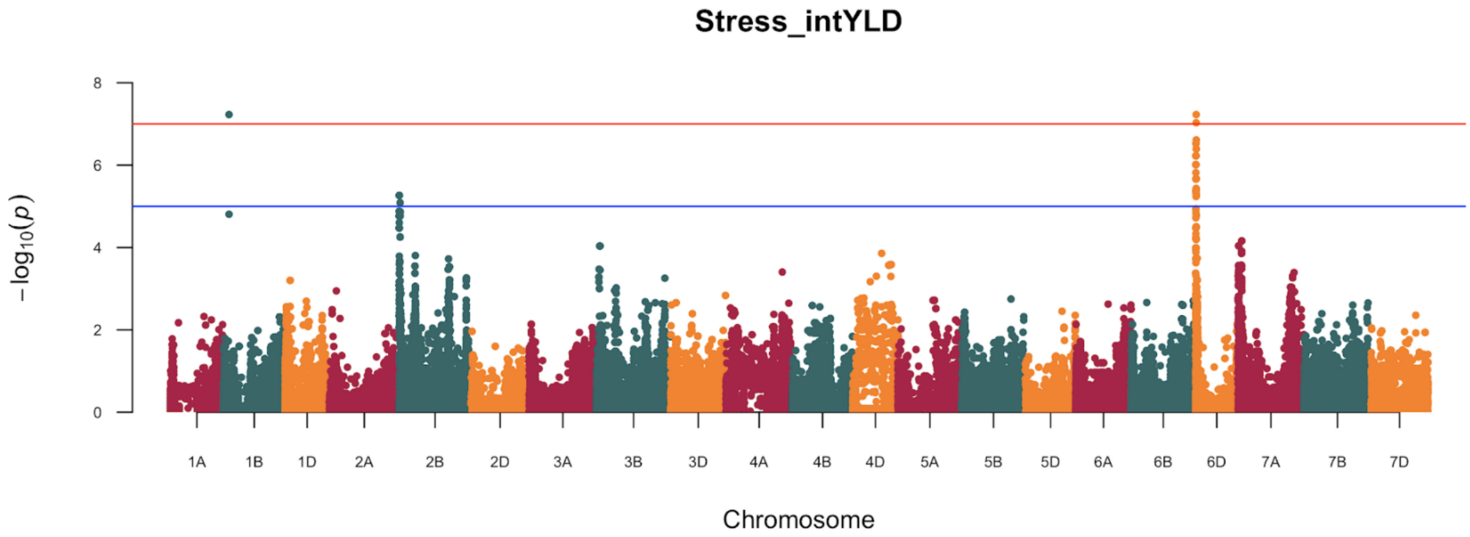
Stress_intYLD
8
6
4
2
0
−log10(p)
1A 1B 1D 2A 2B 2D 3A 3B 3D 4A 4B 4D 5A 5B 5D 6A 6B 6D 7A 7B 7D
Chromosome

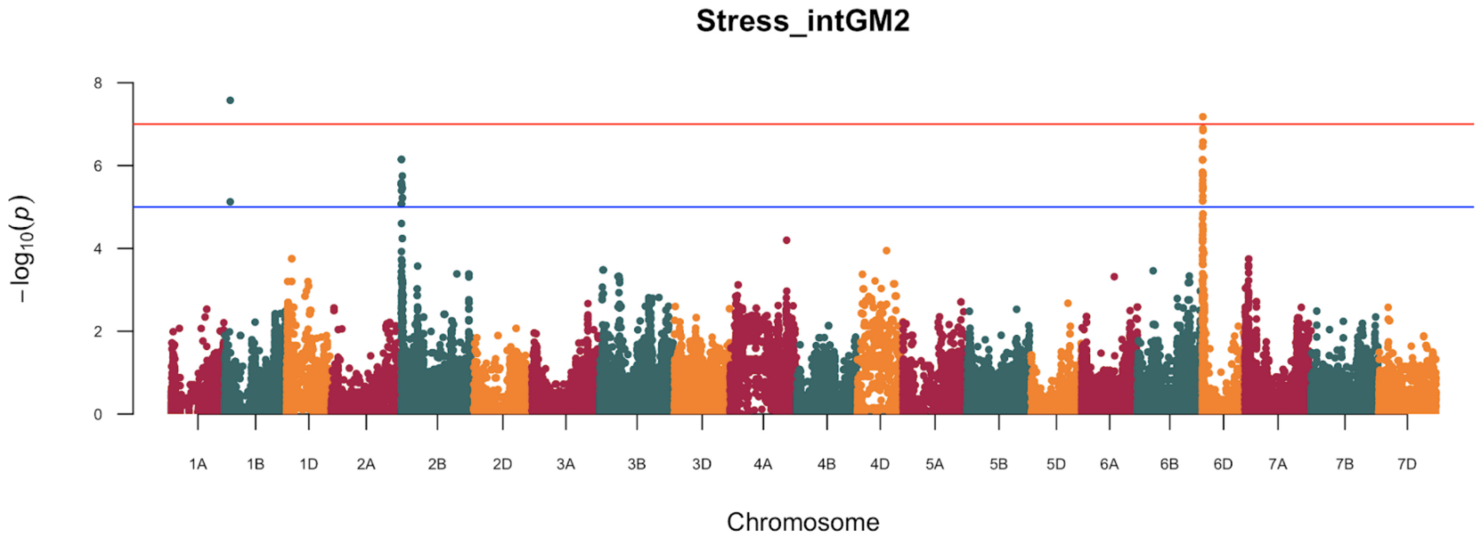
Stress_intGM2
8
6
4
2
0
−log10(p)
1A 1B 1D 2A 2B 2D 3A 3B 3D 4A 4B 4D 5A 5B 5D 6A 6B 6D 7A 7B 7D
Chromosome

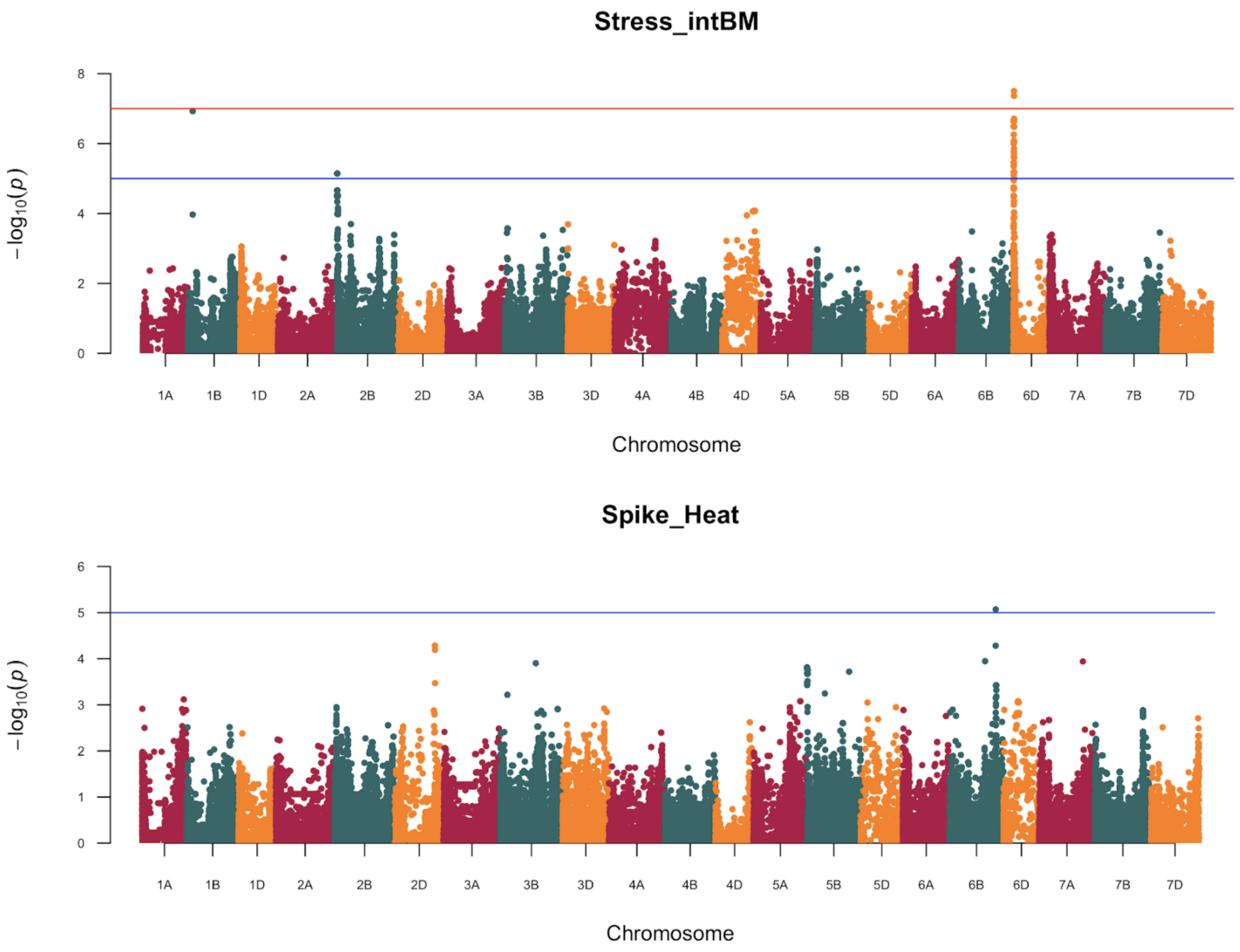
Stress_intBM
−log10(p)
8
6
4
2
0
1A 1B 1D 2A 2B 2D 3A 3B 3D 4A 4B 4D 5A 5B 5D 6A 6B 6D 7A 7B 7D
Chromosome
Spike_Heat
−log10(p)
6
5
4
3
2
1
0
1A 1B 1D 2A 2B 2D 3A 3B 3D 4A 4B 4D 5A 5B 5D 6A 6B 6D 7A 7B 7D
Chromosome

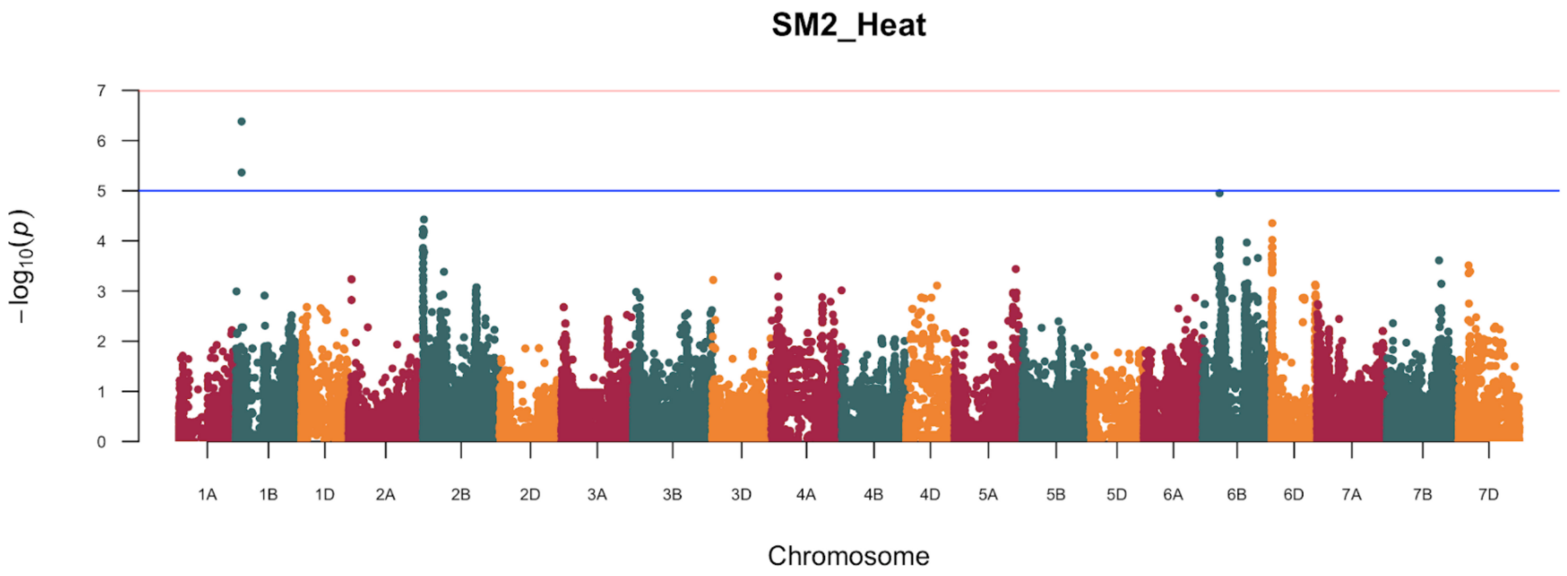
SM2_Heat
−log10(p)
Chromosome
1A 1B 1D 2A 2B 2D 3A 3B 3D 4A 4B 4D 5A 5B 5D 6A 6B 6D 7A 7B 7D

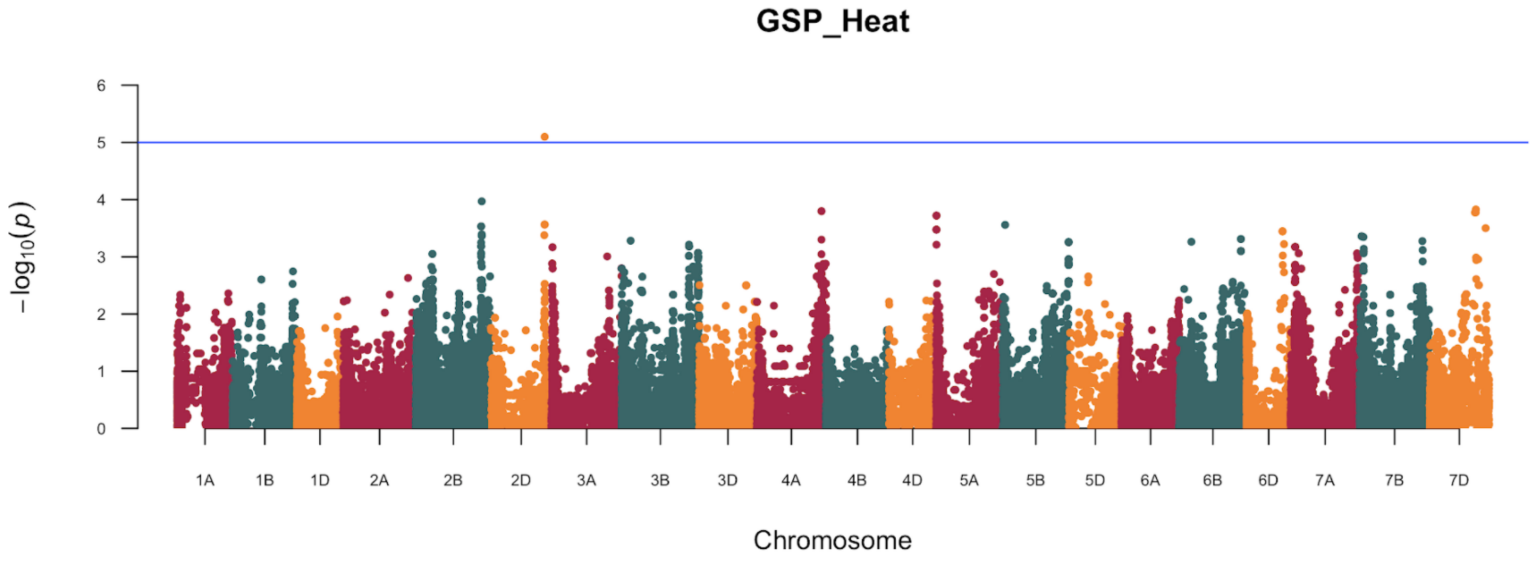
GSP_Heat
−log10(p)
Chromosome
1A 1B 1D 2A 2B 2D 3A 3B 3D 4A 4B 4D 5A 5B 5D 6A 6B 6D 7A 7B 7D

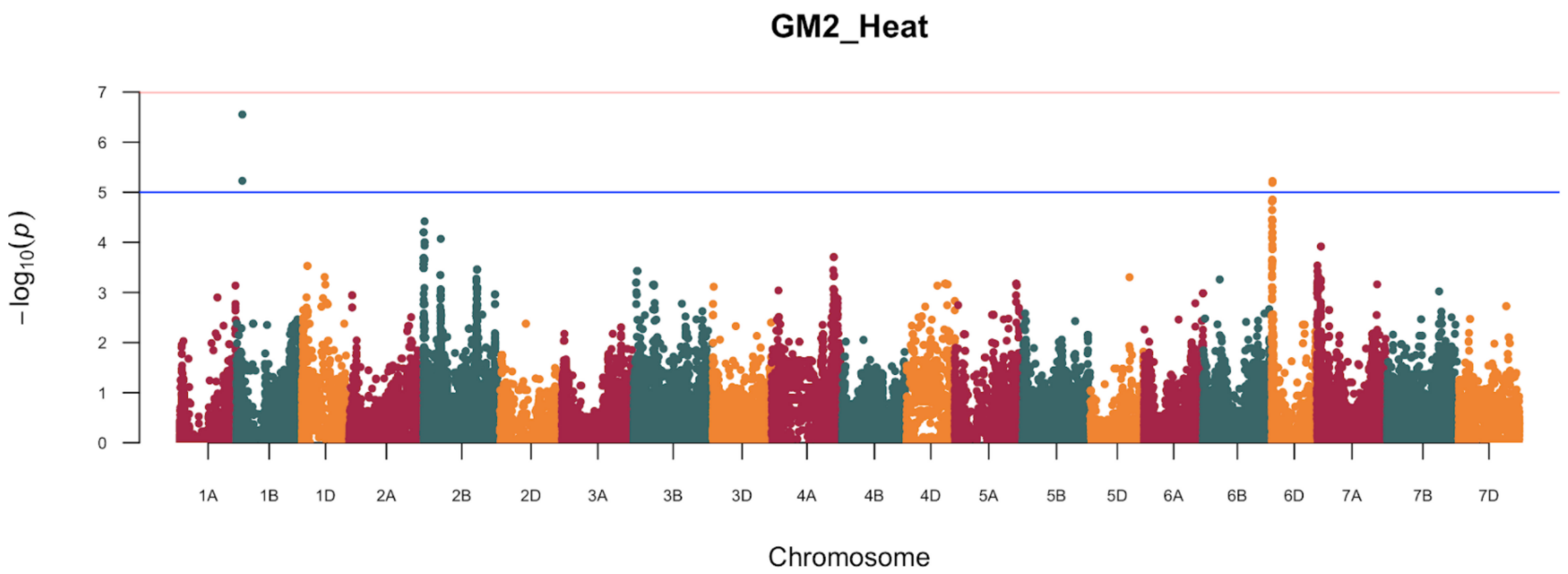
GM2_Heat
$-\log_{10}(p)$
0
1
2
3
4
5
6
7
1A 1B 1D 2A 2B 2D 3A 3B 3D 4A 4B 4D 5A 5B 5D 6A 6B 6D 7A 7B 7D
Chromosome

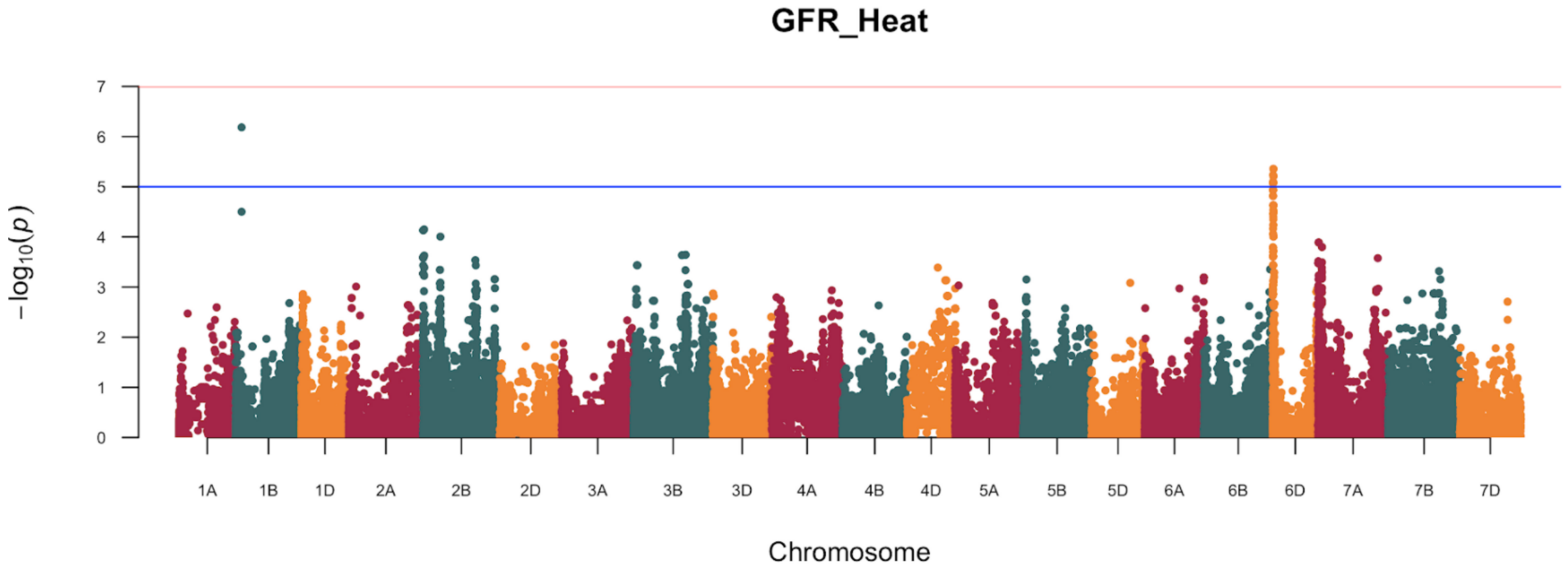
GFR_Heat
$-\log_{10}(p)$
0
1
2
3
4
5
6
7
1A 1B 1D 2A 2B 2D 3A 3B 3D 4A 4B 4D 5A 5B 5D 6A 6B 6D 7A 7B 7D
Chromosome

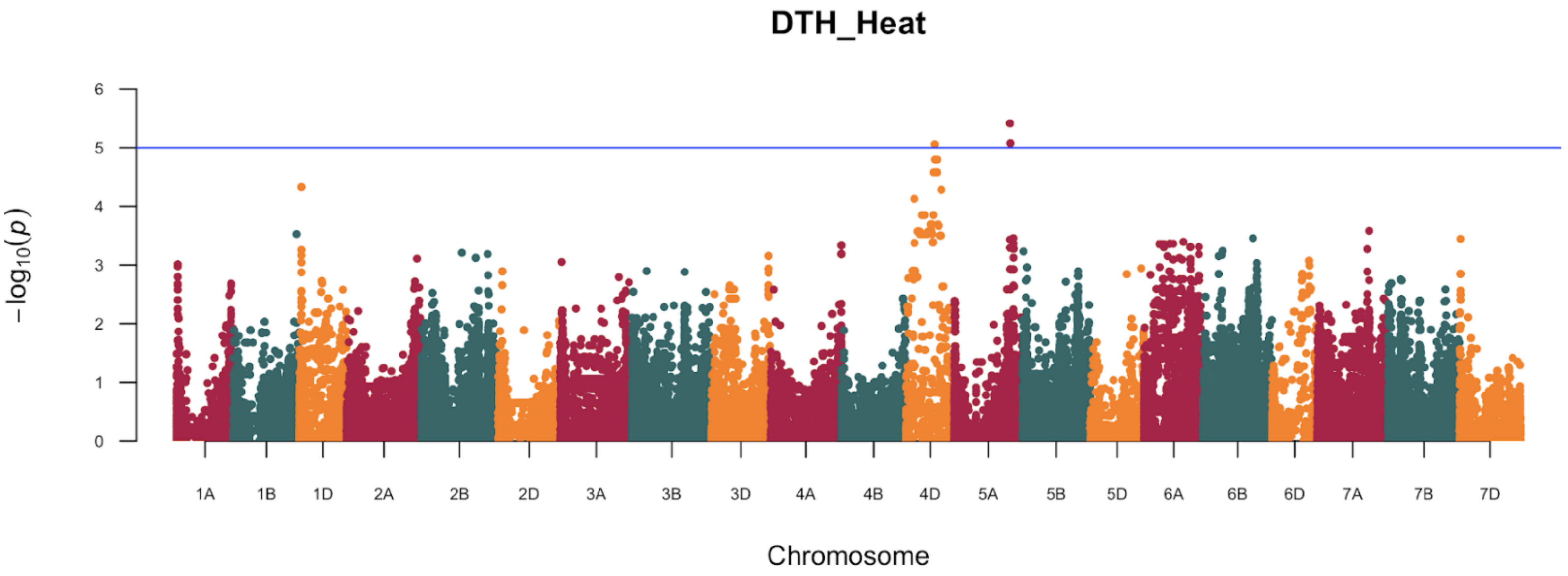


**Supplementary Figure 2.** GWAS results in HiBAP I based on BLUEs means obtained from the combined analysis from Ht16 and Ht17. The dotted horizontal line indicates threshold of significance.

**Figure S3**

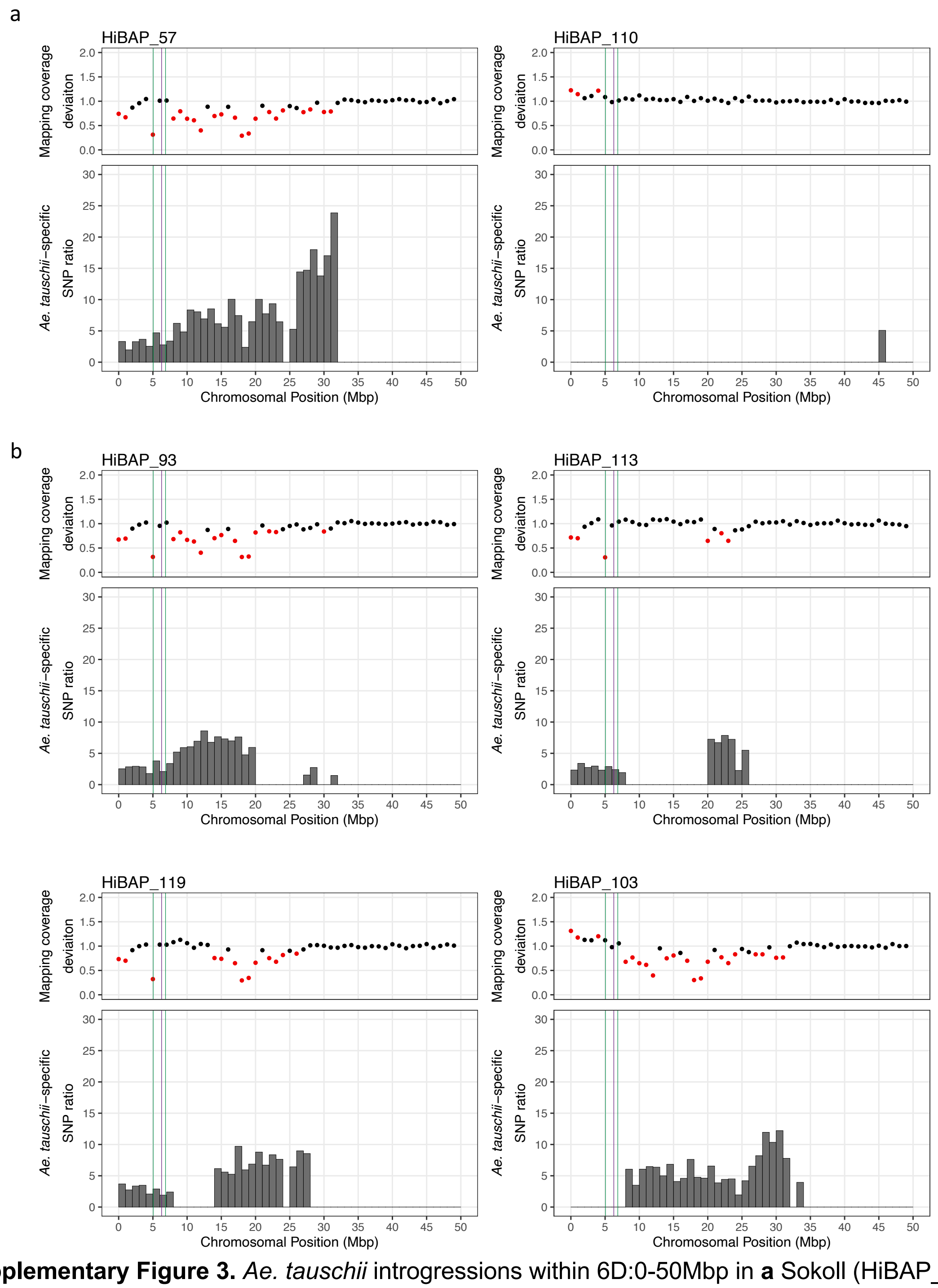


**Supplementary Figure 3.** *Ae. tauschii* introgressions within 6D:0-50Mbp in **a** Sokoll (HiBAP_57) and Weebil1 (HiBAP_110) and **b** four independent Sokoll/Weebil1 crosses. Mapping coverage deviation was computed between the HiBAP line and the median of the panel in 1Mbp windows. Red points are statistically significant outliers. *Ae. tauschii*-specific SNP ratio in each 1Mbp window was calculated by dividing the number of homozygous *Am. muticum*-specific SNPs in that window by mean number of homozgyous SNPs in that window across the panel. Green lines mark the borders of the region common to all lines with the T haplotype, corresponding to a 1.80Mbp region in CS and a 1.49Mbp in *Ae. Tauschii.* The purple line indicates the 6D MTA position.
